# Investigating Regional Specificity of Alzheimer’s Disease Pathology in the 5xFAD Mouse Model

**DOI:** 10.1101/2021.07.08.441803

**Authors:** Araba Gyan, Emily D Glass, Tamara Stevenson, Geoffrey G Murphy, Shannon J Moore

## Abstract

Alzheimer’s disease (AD) is a neurodegenerative disease that affects cognition and memory. Mouse models such as 5xFAD, incorporate mutations found in familial or early-onset (EOAD) AD; these mutations increase the production of the toxic AB species that contributes to plaque formation in the brain. However, not all brain regions in the 5xAD animals accumulate plaques to the same degree. For example, the cerebellum appears to be resistant to AB plaque formation, while other regions such as the cortex and subiculum develop copious plaques. The mechanism(s) underlying this regional specificity remains unclear. Thus, we used a fluorescent antibody to APP/AB to quantify the amount of these proteins in several brain regions of interest, including subiculum, cortex, and cerebellum. We found that the cerebellum had less APP/AB than the other regions quantified, which demonstrates a correlation between the levels of APP/AB and plaque formation. Taken together, this suggests that the regional specificity of AD pathology may be a result of different levels of protein expression.

## INTRODUCTION/BACKGROUND

Alzheimer’s disease (AD) is a progressive neurodegenerative disease that destroys memory and eventually affects the ability to perform basic everyday tasks. It is the most common form of dementia, affecting approximately 5.7 million people in the US and estimated to cost the country about $355 billion, with $239 billion in combined Medicare and Medicaid payments (Alzheimer’s Association, 2021). Furthermore, AD is currently ranked as the sixth leading cause of death in the United States (U.S. Department of Health and Human Services, 2019). Despite advances in the understanding of AD pathogenesis, there are still some unresolved mechanistic questions. In addition, there isn’t an efficacious treatment to prevent or manage symptoms of the disease.

There are two main categories of AD: early-onset AD (EOAD) and late-onset AD (LOAD), or sporadic AD. The majority of AD patients have sporadic AD and develop symptoms later in life (>65 years). Approximately 1-5% of patients have EOAD, with the development of symptoms occurring before the age of 65 (Rogaeva, 2002). Those with EOAD have autosomal-dominant mutations in the amyloid precursor protein (APP), presenilin 1(PS1), and/or presenilin 2(PS2) (Hatami, 2017). Mouse models incorporating these genetic mutations can then be used to investigate the mechanisms underlying the progression of AD.

To date, there are numerous transgenic mouse models of AD incorporate mutations in the humanized APP and/or PS1 genes. One of these is the 5xFAD mouse model that expresses five EOAD (also called familial AD (FAD)) mutations; three mutations in APP, (Swedish [K670N/M671L], Florida [I716V], London [V717I] mutations), and two in PS1 (M146L and L286V) (Oakley, 2006). These mutations significantly increase the production of amyloid-beta (A*β*) peptide and plaque deposition by favoring the amyloidogenic pathway (see below).

### Amyloidogenic and Non-amyloidogenic Processing

APP is an integral membrane protein that is expressed in many tissues. APP is metabolized in two different routes i.e. the amyloidogenic and non-amyloidogenic pathways (Fig 1). In the non-amyloidogenic pathway, the cleavage of APP by *α*-secretase in the region that contains A*β* peptide prevents protein aggregation and reduces the formation of plaques. In the amyloidogenic pathway, APP is cleaved by *β*-secretase and *γ*-secretase, which produces A*β* peptide. These A*β* peptides are released, form oligomers and aggregate into plaques.

**Figure1:**
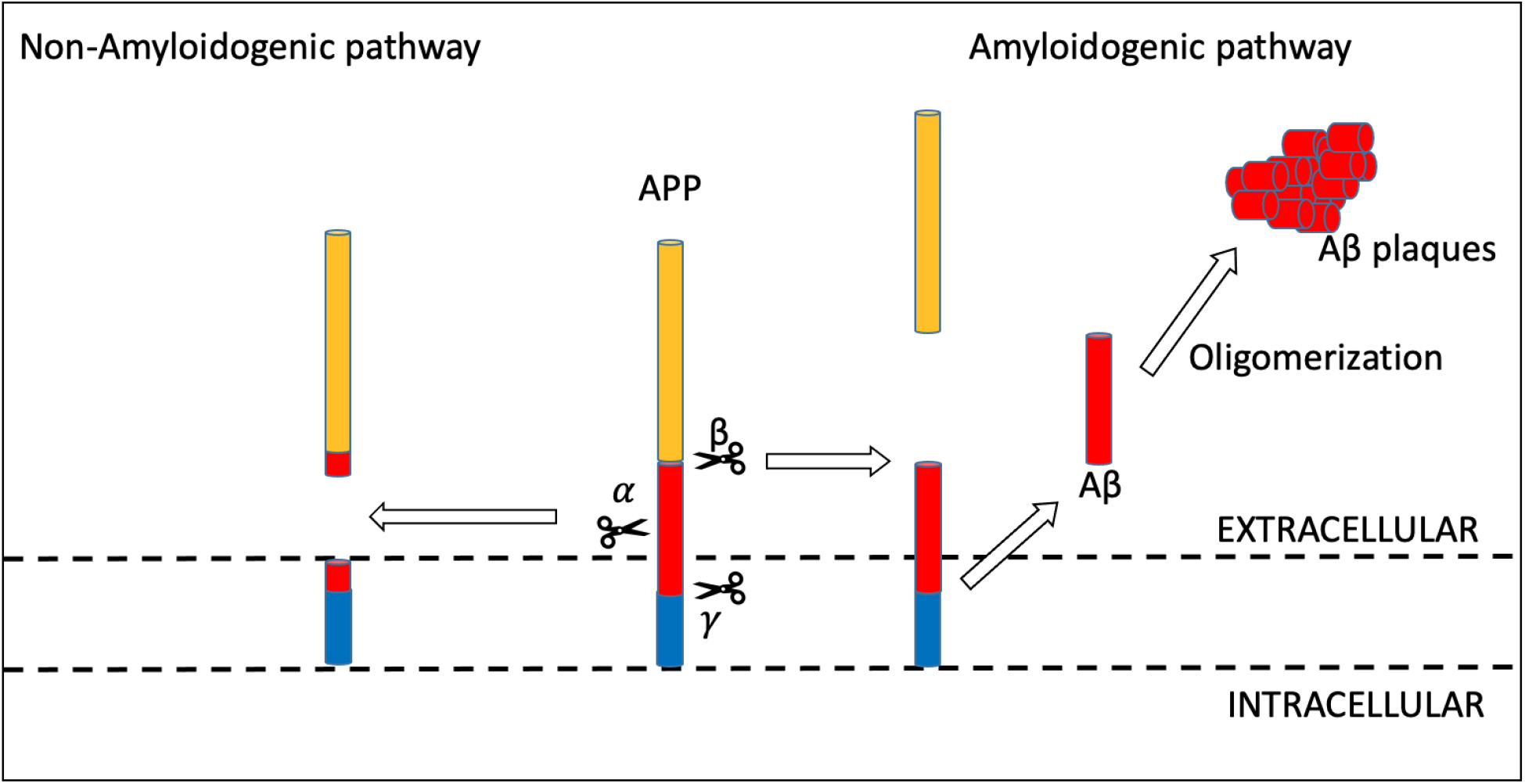
Amyloidogenic and non-amyloidogenic APP-processing pathways. (Right) In the amyloidogenic pathway, amyloid precursor protein (APP) is cleaved by two enzymes; first by β-secretase then *γ*-secretase to produce amyloid-beta (Aβ). These Aβ fragments then aggregate to form plaques. (Left) In the non-amyloidogenic pathway, plaque formation is precluded when *α*-secretase cleaves APP.

Interestingly, not all brain regions seem to accumulate plaques to the same degree. For instance, previous experiments conducted in our lab have shown that some regions of the brain are extremely susceptible to A*β* plaque formation, while other regions appear less vulnerable (Fig 2).

**Figure 2.**
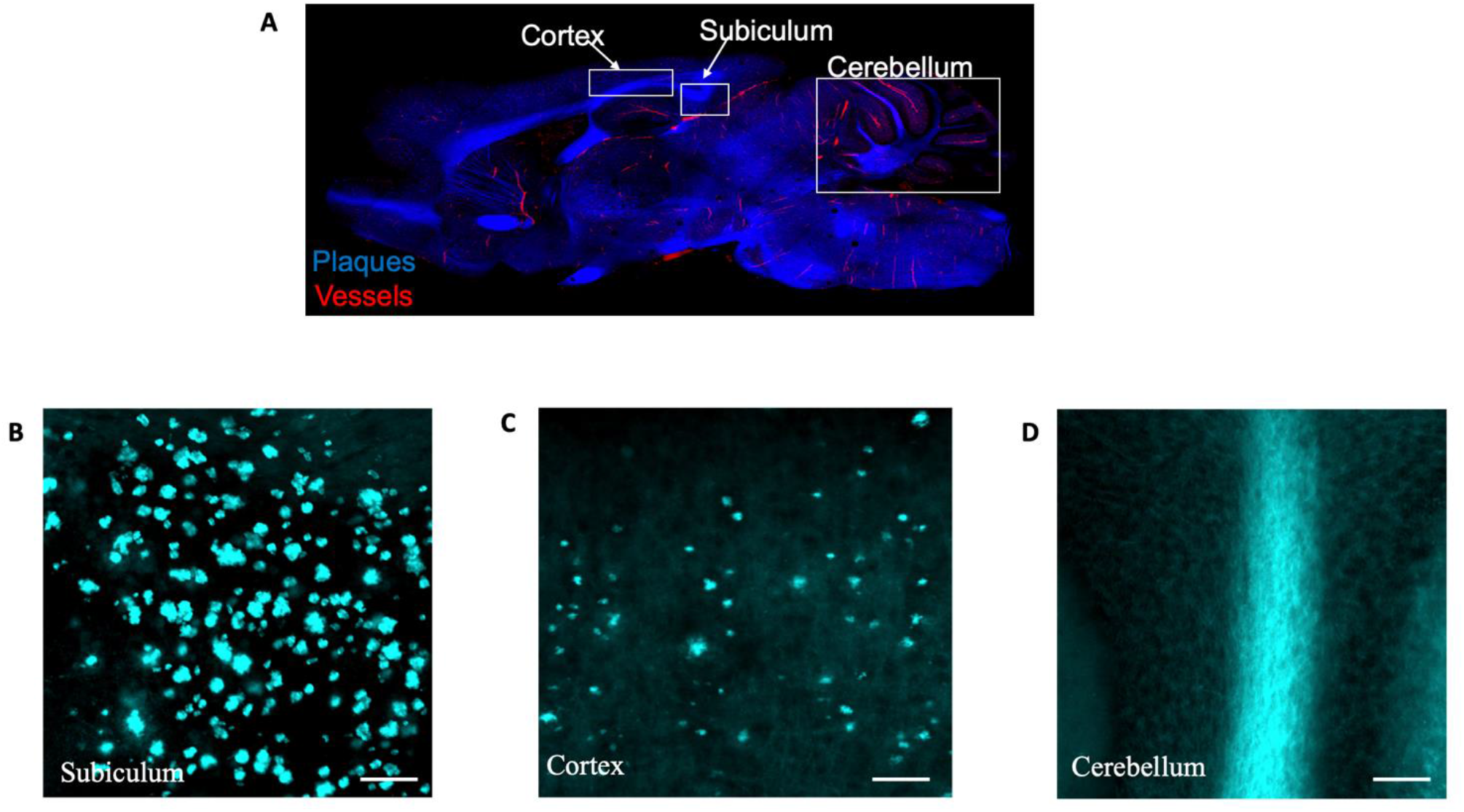
Plaque accumulation in the Subiculum, Cortex and Cerebellum in the 5xFAD mouse model. (A) A tile-scanned image of a brain section with labeled plaques (BTA-1,blue) and blood vessels (tdTomatoe-lectin,red). (B-D) A representative image of three different brain regions to compare the plaque accumulation in each region. Note the presence of plaques in both subiculum (B) and cortex (C), and lack of any plaque deposition in cerebellum (D). Scale bars in B-D, 50um.

For instance, in the 5xFAD mouse model, almost all areas in the brain have extensive plaque accumulation except the cerebellum, which has no apparent plaque accumulation. It is not known why the cerebellum seems to be relatively spared from the development of plaques. This paper aims to examine potential mechanistic regional differences in APP/Aβ levels that could be a contributing factor to the regional specificity observed in A*β* plaque accumulation.

## MATERIALS AND METHODS

### Animals

The 5xFAD transgenic mice, which will be referred to from now on as transgene-positive, were acquired from the Jackson lab on a C57BL/6J background (stock# 34848). 5xFAD mice were then backcrossed onto a C57BL/6Taconic for at least 10 generations by our lab before use for experiments. Experimental animals were approximately 6-8-month-old with approximately equal numbers of male and females and included both transgene-positive and transgene-negative mice. The transgene-negative littermates were used as a control throughout the experiment. All the animals were housed in a controlled suitable environment with 14/10-hour light/dark cycle. They were provided with water and food *ad libitum*. All the animal experiments were approved by the Institutional Animal Care & Use Committee (IACUC) at the University of Michigan and were performed in accordance with the rules and guidelines of the United States Public Health Services Policy on Humane Care and Use of Laboratory Animals.

### Sample Preparations

Mice were put under anesthesia with isoflurane and sacrificed by transcardiac perfusion; Phosphate-buffered Saline (PBS) was perfused for approximately 3 minutes, followed by perfusion with 4% paraformaldehyde (PFA) for approximately 5 minutes. The tissue was then fixed in PFA overnight and cryopreserved using a hypertonic solution of 30% sucrose until the brains sank (2-4 days). The brains were frozen in an Optimal Cutting Temperature compound (OCT) for about an hour on dry ice, then stored in a −80°C freezer until used for sectioning. To section, a brain was attached to the chuck of the cryostat (Leica CM 1860) and sliced sagittally into 40μm sections. Tissues were adhered to Shandon Colorfrost™Plus Microscope Slides (Ref#9991001, Lot#102120-9) and stored in a −80°C freezer until ready for immunolabeling.

### Immunofluorescence labeling

Sections were stained using a common standard immunofluorescence protocol (Frediksson et al., 2015). Additionally, sections underwent antigen retrieval (DAKO, S1700) and permeabilization with 0.5% Triton X-100. The sections were then incubated with primary antibody that recognizes an epitope within the humanized Aβ sequence using Purified Anti-B-Amyloid (BioLegend, Cat#803012, Lot#B251533) directly conjugated with horseradish peroxide (HRP) at a 1:100 dilution ratio and stored at 4 °C overnight. Note that this antibody will label full-length APP before Aβ is released by enzymatic cleavage, and the Aβ fragements/plaques after cleavage. An amplification protocol was also used (Alexa Fluor™488 Tyramide SuperBoost™ kit; Invitrogen by ThermoFisher Scientific, Cat:#B40932, Lot#2169502), following manufacturer’s instructions. The sections were then mounted using VectaShield Antifade Mounting Medium (Vector Laboratories, Cat#H-1000, Lot#ZG0709), coverslipped using Fisherbrand microscope cover glass (ThermoFisher Scientific, Cat#12-545E, Lot#18919) and stored in a dry dark environment until ready to be imaged.

### Image Acquisition

Images were acquired on an inverted Olympus IX83 microscope as non-z-stack images using the Thermo Fisher CellSensor Cell Lines software. The fluorophore was excited using a 488nM wavelength with an exposure time of 50.05ms; images were captured using a 10x objective. For each of the three animals, images were acquired from 3-4 sections of the brain (n=12-13 sections total).

### Quantifìcation/Statistical analysis

All image analysis was performed using FIJI software; images were imported using the BioFormats Importer plugin. Three regions of interest were quantified from each slice: the subiculum, cortex just dorsal to the hippocampus, and the cerebellum. Regions of interest (ROI) were drawn for each brain area and applied to each section. Using the Analyze > Measure function in FIJI, the mean average of the fluorescence intensity of each ROI was measured. Statistical analysis was done using a one-way ANOVA in GraphPad Prism 9 (GraphPad Software, La Jolla, CA, USA) with Tukey’s post hoc test to determine pairwise significance using a significance level of p < 0.05.

## RESULTS

### There is less APP/Aβ in the cerebellum compared to the subiculum and cortex

To investigate why the 5xFAD mouse model doesn’t develop plaques in the cerebellum, we measured humanized APP/A*β* using a fluorescent antibody in different brain regions (Fig 3). Quantification of the fluorescence intensity was used as an indirect measure of the APP/A*β* in the individual brain regions. We assumed no difference between the transgene-negative and transgene-positive animals in either the biological background or the imaging artifacts background. Thus, we decided to use the transgene-negative animals (that should not have humanized APP/A*β* fluorescence) as a control for the biological and imaging artifacts background. We calculated the average of the mean fluorescence intensity of all the transgene-negative animals for each ROI. The mean fluorescence intensity for each transgene-positive animal was then normalized to the respective ROI average from the transgene-negative mice.

**Figure.3.**
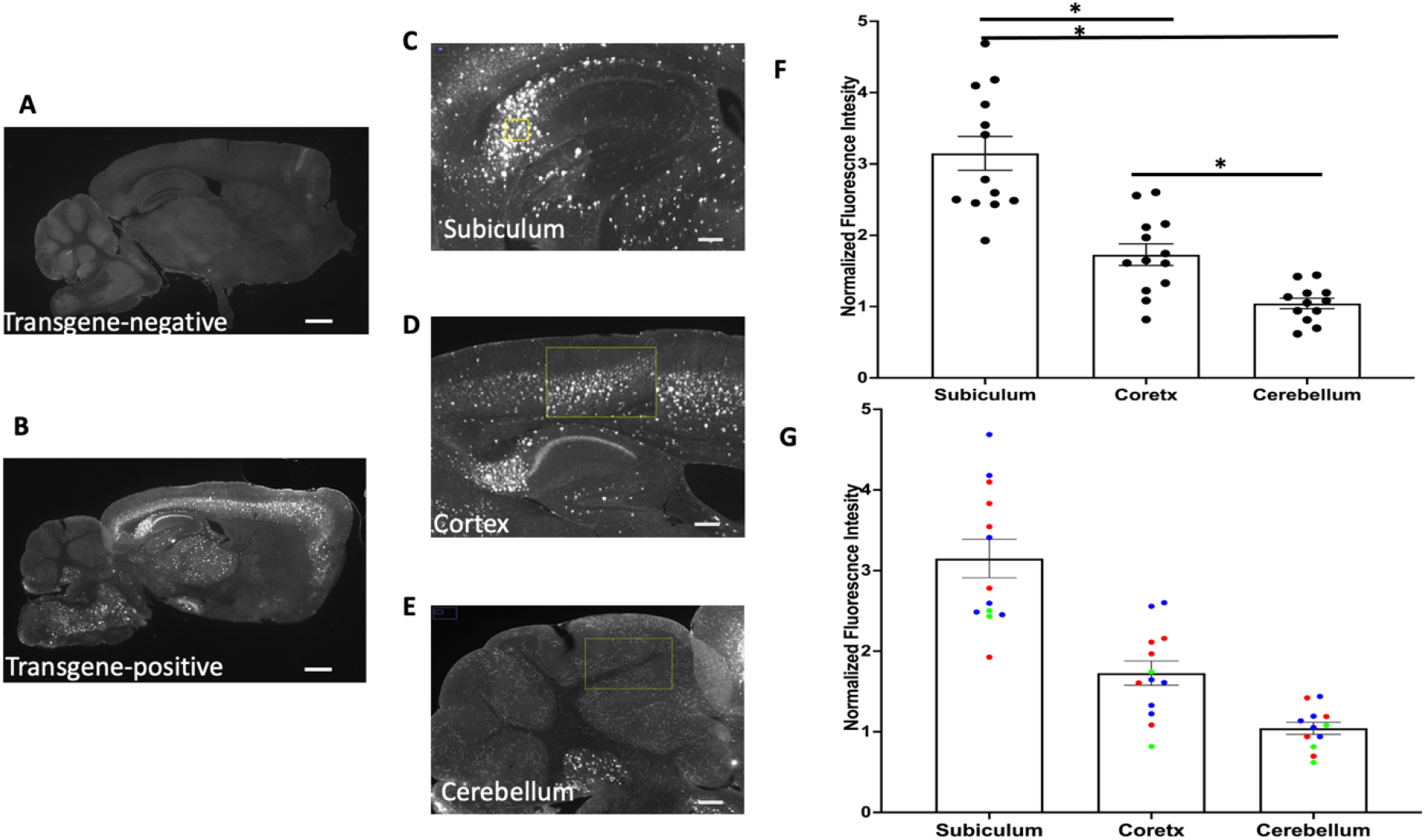
Lower levels of APP/ Aβ in the cerebellum. (A and B) Brain slices from transgene-negative and positive animals stained with purified Anti-beta-Amyloid antibody. Scale bars in A and B, 1000um. (C-E) Representative sections of the brain regions measured for fluorescence intensity. Scale bars in C-E, 50um. (F and G) Quantifications of fluorescence intensity in each region. (F) Fluorescence intensity was significantly higher in subiculum compared to both cortex (p<0.0001) and cerebellum (p<0.0001), and likewise was significantly higher in cortex compared to cerebellum (p=0.0222). (G) The same data as in (F) but color-coded so all sections from the same animal are the same color. Data are presented as mean +/- SEM and analyzed by one-way ANOVA with post hoc Tukey’s test; significance was determined as p<0.05.

We looked at the following regions: subiculum, cortex, and cerebellum. We see an especially strong signal in the subiculum and cortex of the transgene-positive animals. When we analyzed the mean fluorescence intensity, there was a significant difference across regions (one-way ANOVA: F(2,35) = 38.91; p<0.0001; Fig 3F). Using Tukey’s post hoc test, we found that the level of fluorescence in the subiculum was approximately 3.1x that of the cerebellum (p<0.0001), and the level of fluorescence in the cortex was approximately 1.7x compared to the cerebellum (p=0.0222). In addition, subiculum also had an approximately 1.8x higher level of fluorescence compared to cortex (p<0.0001).

## DISCUSSION

In the 5xFAD mouse model, the cerebellum has no plaques. In other mouse models, the cerebellum may have some plaques, but still far fewer than other regions. Hence, the cerebellum seems to be protected. Our data showed that the cerebellum had significantly less APP/A*β* than the other regions quantified, and also has fewer plaques. On the other hand, the subiculum had the highest APP/A*β* levels, and also has more plaques than the cortex and the cerebellum. Thus, APP/A*β* levels appear to correlate with plaque deposition. Less APP/A*β* in the cerebellum may be one reason this region has no plaques in the 5xFAD mouse model. Taken together, less APP/A*β* may be a mechanism that accounts for regional differences in vulnerability to AD pathology.

Besides the mechanism explored in this paper, there are other factors that may also contribute to the very low (to no) plaque deposition observed in the cerebellum. One of these potential mechanisms is the abundance and activity of the enzymes in both the amyloidogenic and non-amyloidogenic pathways. Plaques are generated sequentially after *β*- and *γ*-secretase cleaves the APP in the amyloidogenic pathway. On other hand, cleavage by *α*-secretase (in the non-amyloidogenic pathway) does not lead to plaque formation. Thus, if *α*-secretase is more abundant and/or active in the cerebellum, this may contribute to lower plaque load. Likewise, if the *β*- and *γ*-secretase are less active than in other regions, this would also lead to a lower plaque burden in the cerebellum. Another possibility is that these enzymes have the same activity as in other regions, but that the immune response is more active or efficient in cerebellum. In this case, Aβ is being cleaved, oligomerizing, and aggregating to form plaques at the same rate as in other regions. However, microglia and astrocytes may more efficiently target these plaques for degradation, therefore keeping the cerebellum relatively plaque-free.

Taken together, this study has identified one of the potential mechanisms by which the cerebellum maintains protection against plaque deposition in the 5xFAD. The 5xFAD has no plaques in the cerebellum, and other mouse models also have fewer plaques in the cerebellum than other regions. To answer why the cerebellum seems protected, future research is crucial to gain detailed knowledge of any additional mechanisms contributing to the resilience of the cerebellum to plaque accumulation. This will provide a better understanding of the A*β* pathology of the cerebellum and resilience of this brain region.

